# Disease-Associated Mutations in Human BICD2 Hyperactivate Motility of Dynein-Dynactin

**DOI:** 10.1101/121400

**Authors:** Walter Huynh, Ronald D. Vale

## Abstract

Bicaudal D2 (BICD2) joins dynein with dynactin into a ternary complex (termed DDB) capable of processive movement. Point mutations in the *BICD2* gene have been identified in patients with a dominant form of spinal muscular atrophy, but how these mutations cause disease is unknown. To investigate this question, we have developed in vitro motility assays with purified DDB and BICD2’s membrane vesicle partner, the GTPase Rab6a. Rab6a-GTP, either in solution or bound to artificial liposomes, released BICD2 from an autoinhibited state and promoted robust dynein-dynactin transport. In these assays, BICD2 mutants showed an enhanced ability to form motile DDB complexes. Increased retrograde transport by BICD2 mutants also was observed in cells using an inducible organelle transport assay. When overexpressed in rat hippocampal neurons, the hyperactive BICD2 mutants decreased neurite growth. Our results reveal that dominant mutations in BICD2 hyperactivate DDB motility and suggest that an imbalance of minus- versus plus-end-directed microtubule motility in neurons may underlie spinal muscular atrophy.

## Introduction

The retrograde motor, cytoplasmic dynein, is involved in a host of cellular activities, including the trafficking of diverse cargos such as organelles, vesicles, and mRNA as well as mitotic spindle alignment and chromosome positioning (Allan, 2011). In contrast to yeast dynein, which can achieve long-range transport on its own (Reck-Peterson et al., 2006), mammalian dynein must form a tripartite complex with the 1.2 MD dynactin complex and an adapter protein to move processively along microtubules (McKenney et al., 2014; Schlager et al., 2014a).

A number of different adaptor proteins that join dynein and dynactin into an active motile complex have been identified (McKenney et al., 2014; Cianfrocco et al., 2015). For example, Bicaudal D2 (BICD2), one of best studied dynein adaptors, interacts with the small GTPase Rab6, which is found on early endosomes and ER-Golgi vesicles (Matanis et al., 2002). BICD2 is thought to be auto-inhibited through an interaction between its C-terminal coiled-coil domain (CC3) with the N-terminal domain of the protein, which prevents it from associating with its other partners, including dynein (Matanis et al., 2002; Hoogenraad et al., 2003; Liu et al., 2013). Auto-inhibition is thought to be relieved by binding to an effector, such as Rab6a in its GTP-bound form (Liu et al., 2013); however, evidence for this model is lacking.

*BICD2* recently emerged as a genetic locus associated with dominant spinal muscular atrophy (SMA), a genetic disorder characterized by the degeneration of anterior horn cells and leading to eventual muscle atrophy and weakness. (Neveling et al., 2013; Oates et al., 2013; Peeters et al., 2013). The effects that these point mutations have on BICD2’s ability to function as a dynein adaptor are not known, although overexpression of these mutants has been reported to cause Golgi fragmentation and changes in BICD2 localization in cultured cells (Neveling et al., 2013; Peeters et al., 2013). Interestingly, point mutations in the tail region of heavy chain 1 of cytoplasmic dynein have been associated with another form of spinal muscular atrophy, SMA-LED1. These mutations were reported to cause an increase in the affinity between the motor and BICD2 (Peeters et al., 2015) and a decrease in dynein-dynactin processivity (Hoang et al., 2017).

Here, we, using biochemical and single-molecule motility assays, we demonstrate that BICD2 disease mutations cause a gain-of-function in the motility of DDB complexes both in vitro and in cells. In addition, we show that the expression of BICD2 mutants in neurons leads to impaired neurite outgrowth. These results indicate that the SMA BICD2 mutants hyperactivate dynein motility, a gain-of-function consistent with a dominant genetic trait and which could explain the gradual loss of motor neuron function in SMA.

## Results and discussion

### Activation of BICD2 via Rab6a-GTP increases dynein-dynactin binding and motility

Bicaudal D2 is comprised of five coiled-coil domains that can be grouped into three distinct regions (Fig. 1A). The C-terminal region of the protein is the most well-conserved and contains the binding site to different partners localized to different cargos, including the small GTPase Rab6a (Matanis et al., 2002). An N-terminal construct of the protein, which spans residues 25-400 (BICD2_25-400_), was shown to be more effective in binding dynein-dynactin in cell lysate pull-down assays compared to autoinhibited full-length protein (BICD2_FL_) (Splinter et al., 2012; McKenney et al., 2014). We replicated these findings; the difference in dynactin binding was especially striking, exhibiting about a 10-fold reduction for the BICD2_FL_ pull-down compared with BICD2_25-400_, while dynein was ∼2-3 fold lower (Supp. Fig. 1A and 1B).

**Figure 1:**
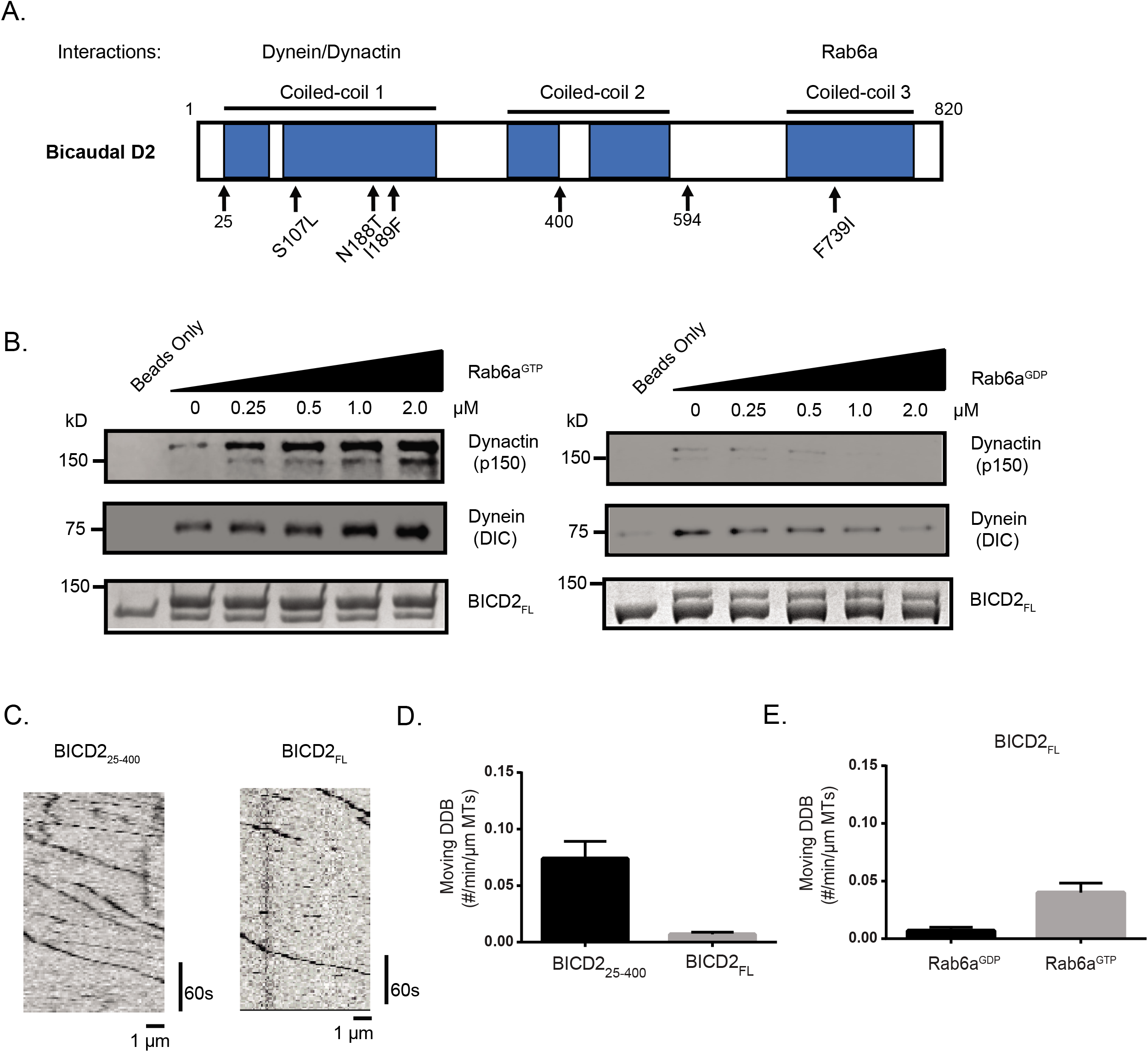
BICD2_FL_ activation by Rab6a^GTP^. **(A)** Domain architecture of mammalian Bicaudal D2. N188T, I189F, and S107L are mutations associated with human SMA-LED, and F739I is the analogous mutation to the classical Drosophila mutation, F684I. Amino acids 400 and 594 indicate the end of constructs used in this study. **(B)** Porcine brain lysate pull-down using full-length BICD2 as bait. Increasing amounts of recombinant Rab6a^GTP^ (left) or Rab6a^GDP^ (right) was added to the lysate, and the amount of endogenous dynein (dynein intermediate chain (DIC) and dynactin (p150 subunit) bound was analyzed by immunoblot. **(C)** Sample kymographs of single molecule DDB (a.a. 25-400 or full-length (FL) BICD2) motility on microtubules. Fluorescence is from the N-terminal sfGFP tag on BICD2. **(D)** Quantification of the number of moving motors per micron of microtubule; mean and SEM from n = 3 independent experiments (each experiment measuring a minimum of 30 microtubules). **(E)** Quantification of number of moving motors per micron of microtubule of BICD2_FL_ with either Rab6a^GDP^ or Rab6a^GTP^ added; mean and SEM from n = 3 independent experiments (each experiment measuring a minimum of 30 microtubules).

We next tested whether the auto-inhibited state of BICD2_FL_ could be activated by Rab6a. We added purified recombinant Rab6a-Q72L, which is locked in a GTP-bound state (hereon referred to as Rab6a^GTP^), to porcine brain lysate and performed pull-downs using BICD2 as bait (Matanis et al., 2002). With increasing Rab6a^GTP^, there was a corresponding increase of dynein and dynactin in the BICD2_FL_ pull-down. Conversely, the GDP-bound mutant of Rab6a (Rab6a-T27N, hereon Rab6a^GDP^) appeared to somewhat reduce the pull-down of dynein and dynactin (Fig. 1B). Rab6a^GTP^ also was detected in the pellet fraction in these experiments, whereas Rab6a^GDP^ was absent (Supp. Fig. 1C). A densitometric analysis shows a ∼2.5-fold increase in dynein enrichment and a ∼10-fold increase in dynactin at the highest Rab6a^GTP^ concentration used (Supp. Fig. 1D). The greater pull-down of dynactin relative to dynein might reflect an interaction of the Rab6a^GTP^-BicD2 complex with dynactin alone. Together, these results indicate that Rab6a^GTP^ is able to activate full-length BICD2 and enhance its ability to bind to dynein and dynactin.

To further examine whether Rab6a^GTP^ can activate BICD2_FL_ and enable dynein-dynactin motility, we utilized total internal reflection microscopy (TIRF) to visualize DDB motility on microtubules. Purified dynein and dynactin from mammalian RPE-1 cell lysates were incubated with similiar concentrations of either superfolder-GFP (sfGFP)-fused BICD2_25-400_ or BICD2_FL_, and the numbers of moving motor complexes were analyzed via kymographs (Fig. 1C). This single molecule in vitro assay revealed that the truncated BICD2_25-400_ produced more processive dynein-dynactin complexes when compared to BICD2fl, which is again consistent with the idea that BICD2_FL_ is autoinhibited (Fig. 1C, 1D, Supp. Video 1). The addition of Rab6a^GTP^ to BICD2_FL_ and dynein-dynactin resulted in a 3-4-fold increase in the number of moving DDB_FL_ complexes on microtubules compared to Rab6a^GDP^ or the individual DDB components alone (Fig. 1E). In contrast, Rab6a^GTP^ neither increased the motility of DDB_25-400_ (Supp. Fig. 1E) nor interacted with BICD2_25-400_ (Supp. Fig. 1F). Collectively, these results suggest that Rab6a^GTP^ is capable of alleviating the auto-inhibition of full-length BICD2 and activating its ability to form processive DDB complexes.

### Disease-related point mutations in BICD2 increase binding to dynein/dynactin and motility

Next, using our assays, we sought to examine the effects of three BICD2_FL_ mutants identified in patients afflicted with DC-SMA/SMA-LED2 (S107L, N188T, I189F) and a mutant (F739I) that is equivalent to the *Drosophila* F684I mutant that produces a bicaudal phenotype (Fig. 1A) (Wharton and Struhl, 1989; Mohler and Wieschaus, 1986). The latter mutation has been shown to have higher affinity towards dynein as compared to wild-type and was proposed to be more readily activated upon binding to cargo (Liu et al., 2013). All four of the mutated residues are highly conserved in the BICD protein family.

In a brain lysate pull-down assay with Rab6a^GTP^, the four BICD2_FL_ mutants recruited more dynein and dynactin compared to wild-type BICD2_FL_, with the greatest difference being observed in dynactin binding (Fig. 2A, Supp. Fig. 2B). Weak binding of dynein-dynactin also was observed with Rab6a^GDP^ or without Rab6 (Fig. 2A, 2B, Supp. Fig. 2A, 2B). It is possible that endogenous Rab6a in the brain lysate might be responsible for this weak BICD2 mutant-specific pull-down or that other factors in the lysate allow for a loss of autoinhibition. In the single molecule TIRF assay in the presence of Rab6a^GTP^, three of the BICD2_FL_ mutants (N188T, I189F, S107L) showed a statistically significant 20-30% increase in the number of motile DDB compared to wild-type BICD2_FL_ (Fig. 2C). Neither the velocity (Fig. 2D) nor run-length of DDB (Fig. 2E, Supp. Fig. 2C) was different between wild-type and mutant BICD2_FL_. In the presence of Rab6a^GDP^, however, motility was much lower and no statistical difference was observed between mutant and wild-type BICD2_FL_ (Supp. Fig 2D). No significant difference also was observed in the number of motile DDB complexes formed by wild-type and mutant truncated BICD2_25-400_ (Fig. 2F). Wild-type and mutant BICD2_25-400_ also pulled down similar amounts of dynein and dynactin (Supp. Fig. 2E). Collectively, these results indicate the point mutations in BICD2_FL_ enhance Rab6a^GTP^-dependent activation from the auto-inhibited state, producing more motile DDB_FL_ complexes, while not altering the measured parameters of DDB_FL_ motility.

**Figure 2:**
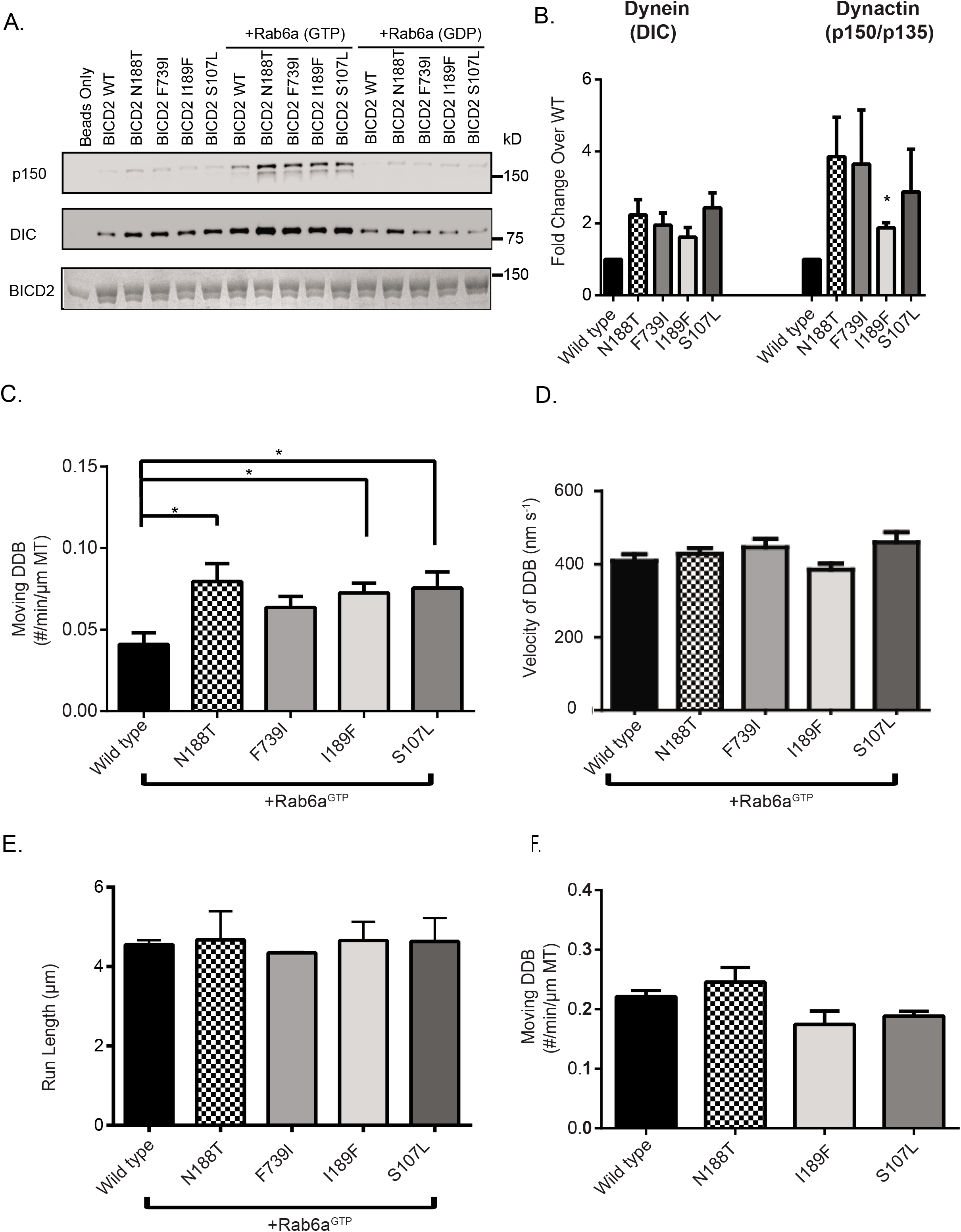
BICD2_FL_ mutants show increased binding to dynein/dynactin and single molecule motility in vitro. **(A)** Porcine brain lysate pull-down using BICD2_FL_ as bait. Rab6a^GTP^ or Rab6a^GDP^ was added to the lysate at a concentration of 1 μM. The amount of endogenous dynein (DIC) and dynactin (p150 subunit) bound was analyzed by immunoblot. The amount of BICD2 on beads was visualized using Coomassie stain. **(B)** The ratio of dynein or dynactin band intensity for mutants compared to wild-type BICD2 for the lysate only condition; mean and SEM from n = 3 independent experiments. Mean, SEM, and p-values for the mutants shown for DIC are as follows: N188T: 2.24 ± 0.42, p=0.10; F739I: 1.95 ± 0.34, p=0.11; I189F: 1.61 ± 0.27, p=0.15; S107L: 2.44 ± 0.41, p=0.07. Mean, SEM, and p-values for p135/p150 are: N188T: 3.86 ± 1.09, p=0.12; F739I: 3.65 ± 1.5, p=0.22; I189F: 1.88 ± 0.14, p=0.027; S107L: 2.88 ± 1.18, p=0.25. Quantitation of the other conditions are shown in Supp. Fig 2A and 2B. **(C)** Dynein and dynactin (purified from RPE-1 cells; see Methods) were incubated with BICD2 and flowed into a motility chamber. The number of moving DDB motors, as visualized by sfGFP fluorescence on BICD2, was quantified per min per micron length of microtubules in the assay; mean and SEM from n = 3 independent experiments. (each experiment measuring a minimum of 30 microtubules) [*P ≤ 0.05]. **(D)** Velocities of the moving motors from part C is shown; mean and SEM from n = 3 independent protein preparations and experiments (each experiment measuring a minimum of 100 DDB complexes). **(E)** The run lengths of WT and mutant DDB_FL_; mean and SEM from n = 3 independent experiments (each experiment measuring a minimum of 130 DDB complexes). For processivity measurements, the NaCl concentration was increased from 50 to 65 mM to reduce the run length so that it could be more reliably measured (see Methods). **(F)** Purified dynein and dynactin were incubated with truncated BICD2_25-400_ (WT and mutant) and flowed into a motility chamber. The number of moving DDB complexes, as visualized by SNAP-TMR fluorescence on BICD2, was quantified per min per micron of microtubule; mean and SEM from n = 3 independent protein preparations and experiments (each experiment measuring a minimum of 30 microtubules).

### A liposome assay provides an in vitro system for cargo transport

The retrograde transport of native membranous cargos (Grigoriev et al., 2007; Utskarpen et al., 2006) involves prenylated, membrane-bound Rab6a interacting with BICD2. We sought to recapitulate this cargo motility in vitro using recombinant Rab6a^GTP^ or Rab6a^GDP^ covalently linked to large unilamellar vesicles (LUVs) via maleimide lipids (Fig. 3A). With Rab6a^GTP^ on the liposomes, we observed the frequent binding and movement of liposomes along microtubules in the presence of dynein, dynactin, and BICD2_FL_ (Fig. 3B, 3C). In contrast, motility events were only rarely observed with Rab6a^GDP^-coated liposomes (Fig. 3B, 3C, Supp. Video 2). Next, we examined the BICD2_FL_ mutants in the Rab6a^GTP^ liposome motility assay. All four mutants produced significantly more moving Rab6a^GTP^ liposomes compared to wild-type BICD2_FL_ (Fig. 3D). The velocities of wild-type and mutant BICD2 were similar (∼500 nm/s; (Supp. Fig. 2F)) and slightly higher than the single molecule velocities. Thus, consistent with the single-molecule motility assays, the BICD2 mutants hyper-activated dynein-dynactin motility in an in vitro cargo transport assay.

**Figure 3:**
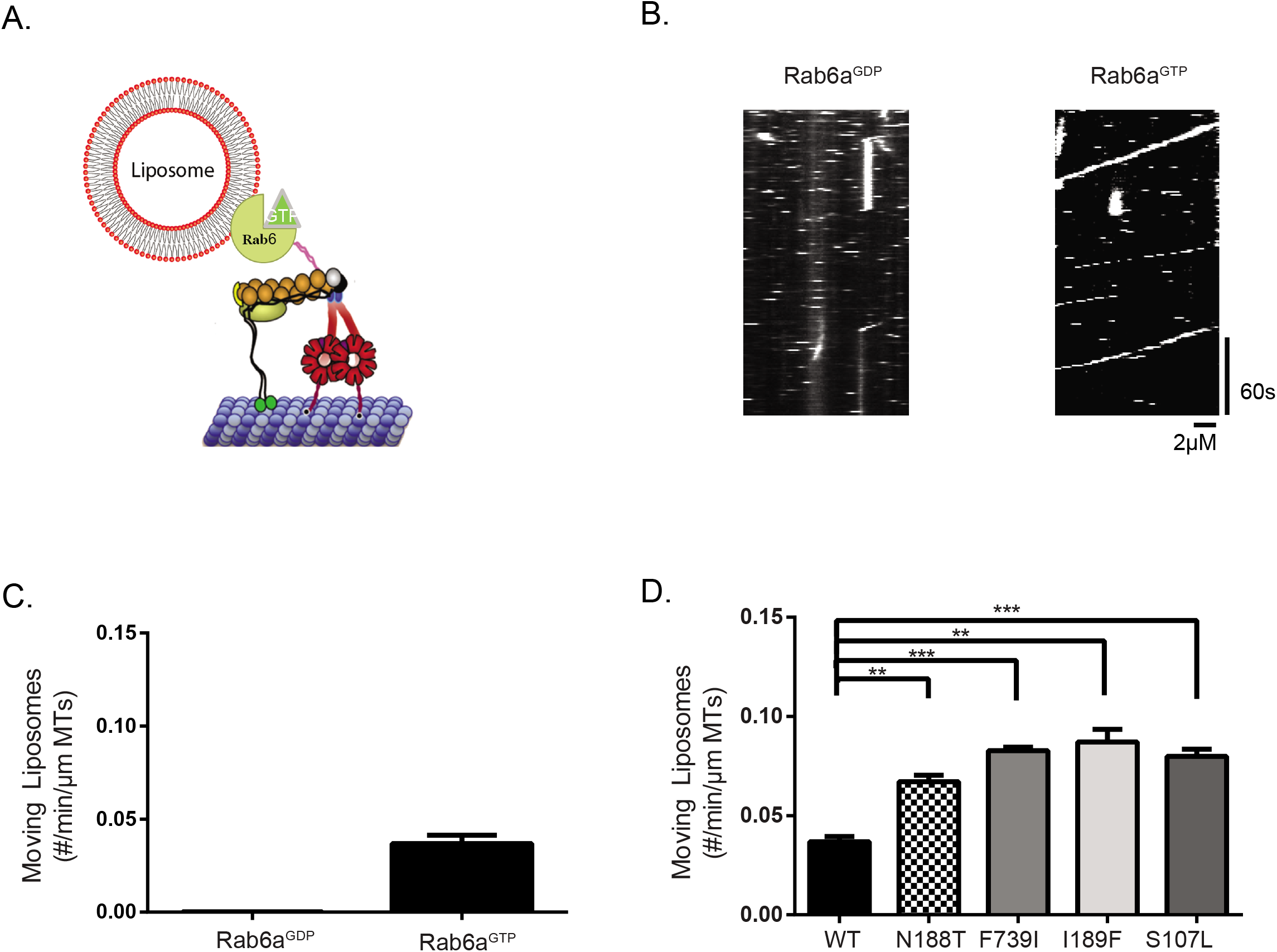
A motility assay using liposomes recapitulates single molecule data. **(A)** A schematic of the liposome motility assay. Rab6a was conjugated to maleimide lipids incorporated into 200 nm-sized liposomes and incubated in solution with DDB. Liposomes were labeled with 0.1% rhodamine-PE. **(B)** Representative kymographs of Rab6a^GDP^ vs Rab6a^GTP^ liposomes moving along microtubule in the presence of DDB_FL_. Horizontal dash lines represent liposomes transiently entering the field of focus. **(C)** Quantification of number of motile Rab6a^GDP^ or Rab6a^GTP^ liposomes when incubated with DDB_FL_; mean and SEM from n = 3 independent experiments (each experiment measuring a minimum of 30 microtubules. Runs for Rab6a^GDP^ occurred very infrequently). **(D)** Quantification of the number of motile Rab6a^GTP^ liposomes moving on microtubules when incubated with DDB_FL_ (wild-type (WT) vs mutant); mean and SEM from n = 3 independent experiments (each experiment measuring a minimum of 30 microtubules) [***P ≤ 0.001, **P ≤ 0.01].

### Enhanced retrograde motility induced by BICD2 mutants in a cell-based assay

To examine the behavior of wild-type and mutant BICD2 in cells, we turned to a previously established inducible cargo trafficking assay (Hoogenraad et al., 2003; Kapitein et al., 2010). In this assay, BICD2-FKBP is co-expressed with FRB fused to a peroxisome localization sequence (Fig. 4A). As shown previously, an N-terminal construct of BICD2 (BICD2_1-594_-FKBP) in the presence of rapamycin increased retrograde transport of peroxisomes and produced significant clustering of the GFP-peroxisome signal around the perinuclear region (Hoogenraad et al., 2003) (Fig. 4B). This longer construct includes part of coiled coil 2, but still lacks the C-terminal region needed for autoinhibition. In contrast, the autoinhibited BICD2_FL_-FKBP construct displayed a more dispersed peroxisome signal (Fig. 4B).

**Figure 4:**
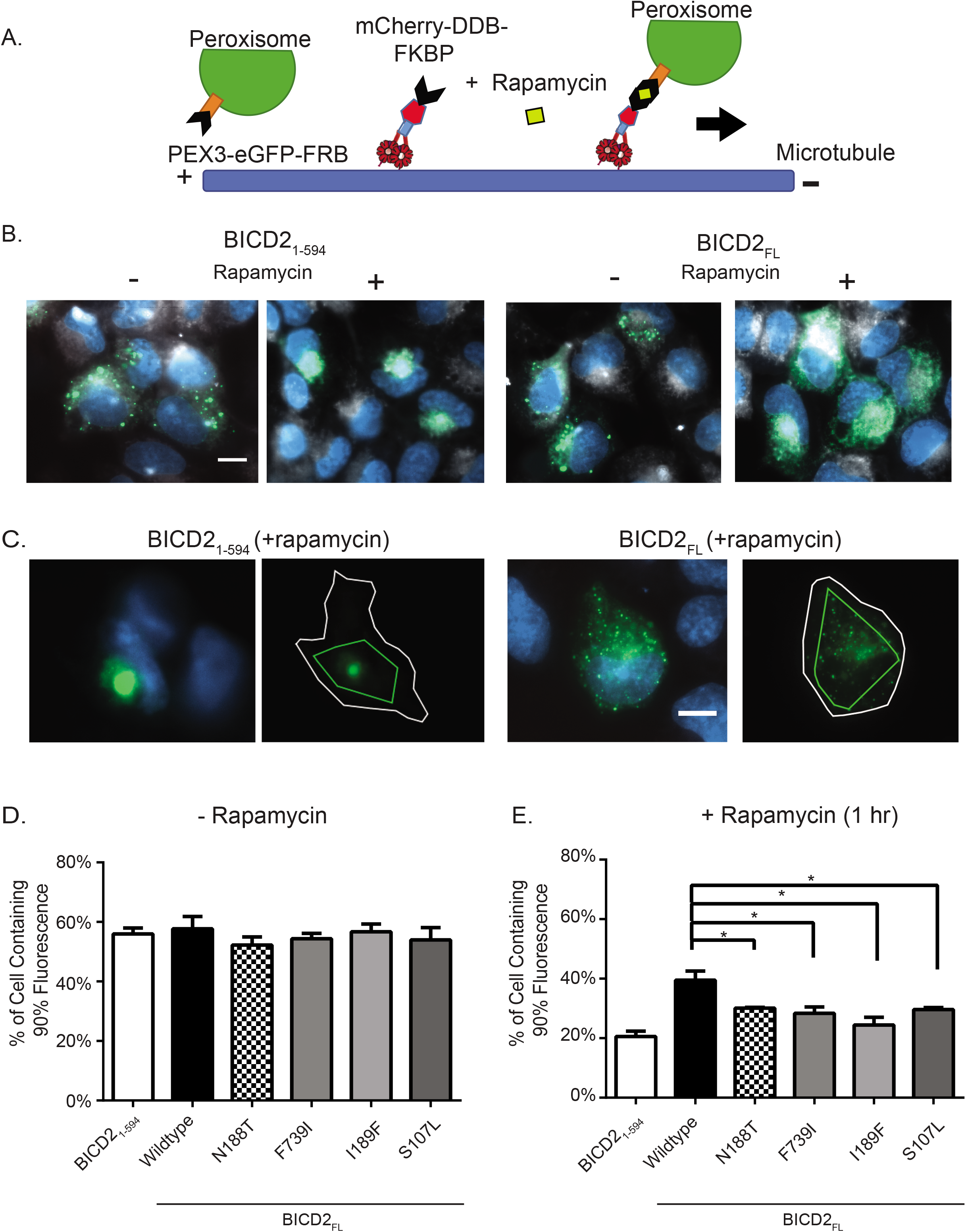
Cellular dynein-dynactin motility induced by wild-type and mutant BICD2. **(A)** Schematic of the cell-based peroxisome motility assay. U2OS cells were co-transfected with a plasmid expressing GFP-FRB with a PEX3 localization sequence, targeting the protein to peroxisomes, and a mCherry-BICD2-FKBP construct. After addition of rapamycin, mCherry-BICD2-FKBP is recruited to peroxisomes. If in an active state, the BICD2 recruits dynein-dynactin which transport the peroxisome along microtubules towards the centrosome. **(B)** Representative epi-fluorescence images of the assay for both BICD2_1-594_ and BICD2_FL_. Perixosomes are shown in green, the white signal represents the far-red cell membrane marker, CellMask, and blue is DAPI. In the absence of rapamycin, peroxisomes decorated with GFP-FRB are distributed throughout the cytoplasm (columns 1 and 3). 1 hr after rapamycin addition, peroxisomes in cells expressing BICD2_1-594_ are heavily clustered at the cell center (column 2), while those expressing BICD2_FL_ exhibit less clustering (column 4). Scale Bar: 10 μm. **(C)** Example image of an analyzed cell for both BICD2_1-594_ and BICD2_FL_. The white outline depicts the cell boundary; the inner green line indicates the area that contains 90% of fluorescence (see Methods). Scale Bar: 10 μm. **(D)** and **(E)** Quantification of the cell clustering data. The percentage of the cell area in which 90% of total fluorescence was contained is used as a measure for the degree of clustering (see Methods). Lower values reflect a higher degree of clustering; mean and SD from n = 3 independent experiments (each experiment measuring a minimum of 25 cells for each construct) [*P ≤ 0.05].

We next tested the mutant BICD2_FL_ constructs in this inducible peroxisome motility assay. To quantify the degree of peroxisome clustering due to minus-end-directed motility, we used a previously created plugin for the image analysis software ICY that calculates the fraction of the cell area containing 90% of the total peroxisome GFP signal; increased transport to the centrosome results in a more clustered signal and thus 90% of the signal occupies less area (de Chaumont et al., 2012; Mounier et al., 2012) (Fig. 4C). In the absence of rapamycin, the clustering values for the wild-type and mutants were similar (∼55%; Fig. 4D). With the addition of rapamycin to cells expressing BICD2_1-594_, the peroxisomes occupied ∼20% of the cell area compared with ∼40% expressing auto-inhibited BICD2_FL_. The four mutants tested in the BICD2_FL_ construct, while not achieving the same degree of peroxisome clustering observed for BICD2_1-594_, showed an increase in the compaction of the peroxisome signal compared to wild-type BICD2_FL_ (Fig. 4D). Thus, similar to the in vitro data, the mutant BICD2_FL_ proteins hyperactivated dynein-dynactin in a cellular context and more efficiently transported peroxisomes towards the centrosomes.

We also tested whether localizing Rab6a^GTP^ containing a peroxisome targeting sequence (Rab6a^GTP^-PEX) could increase the activity of FKBP-BICD2_FL_. However, we found that Rab6a^GTP^-PEX alone (without FKBP-BICD2_FL_ co-expression) was sufficient to cluster peroxisomes, presumably through the recruitment of endogenous BICD2 and then dynein-dynactin (Supp. Fig. 3A, 3B, 3C). Thus, targeting of Rab6a^GTP^ on peroxisome membranes is sufficient for the recruitment and activation of endogenous BICD2 to produce dynein-mediated transport of peroxisomes

### Over-expression of BICD2 mutants in hippocampal neurons decreases neurite length

We next examined whether the hyperactivating BICD2_FL_ mutants affected neurite outgrowth and neuronal morphology. Rat hippocampal neurons, dissected from embryonic day 18 tissue and grown in culture, were transfected with either wild-type or mutant mouse mCherry-BICD2_FL_ constructs along with soluble GFP to visualize the cell body and neurites (Fig. 5A). Three days later, dual mCherry and GFP positive neurons were then scored for the length of their longest process as well as the total length of all processes. Cells transfected with wild-type mCherry-BICD2_FL_ and GFP showed similar neurite lengths compared to cells transfected with GFP alone (Fig. 5B, 5C). This result is consistent with previous work showing that over-expression of wildtype BICD2 in either hippocampal or rat DRG neurons does not affect axon length or overall neurite length (Schlager et al., 2014b). However, cells transfected with the disease-mutant BICD2_FL_ constructs for three days showed a ∼40% decrease in both the length of the longest neurite as well as an overall decrease in total neurite length (Fig. 5B, 5C). This result is similar to what was reported when the BICD-related protein, BICDR-1, which forms a higher velocity DDB complex, is over-expressed in neurons. (M. A. Schlager, Serra-Marques, et al., 2014; M. A. Schlager et al., 2010). The number of long (>15 μm) processes emanating from the cell body was not affected by the BICD2 mutants (Supp. Fig. 3D). The Drosophila mutant (F739I) also showed a significant reduction in total neurites, but not in the axon length (Supp. Fig. 3E, 3F). These results demonstrate that mutations in BICD2 affect the process of neurite outgrowth of hippocampal neurons in culture.

**Figure 5:**
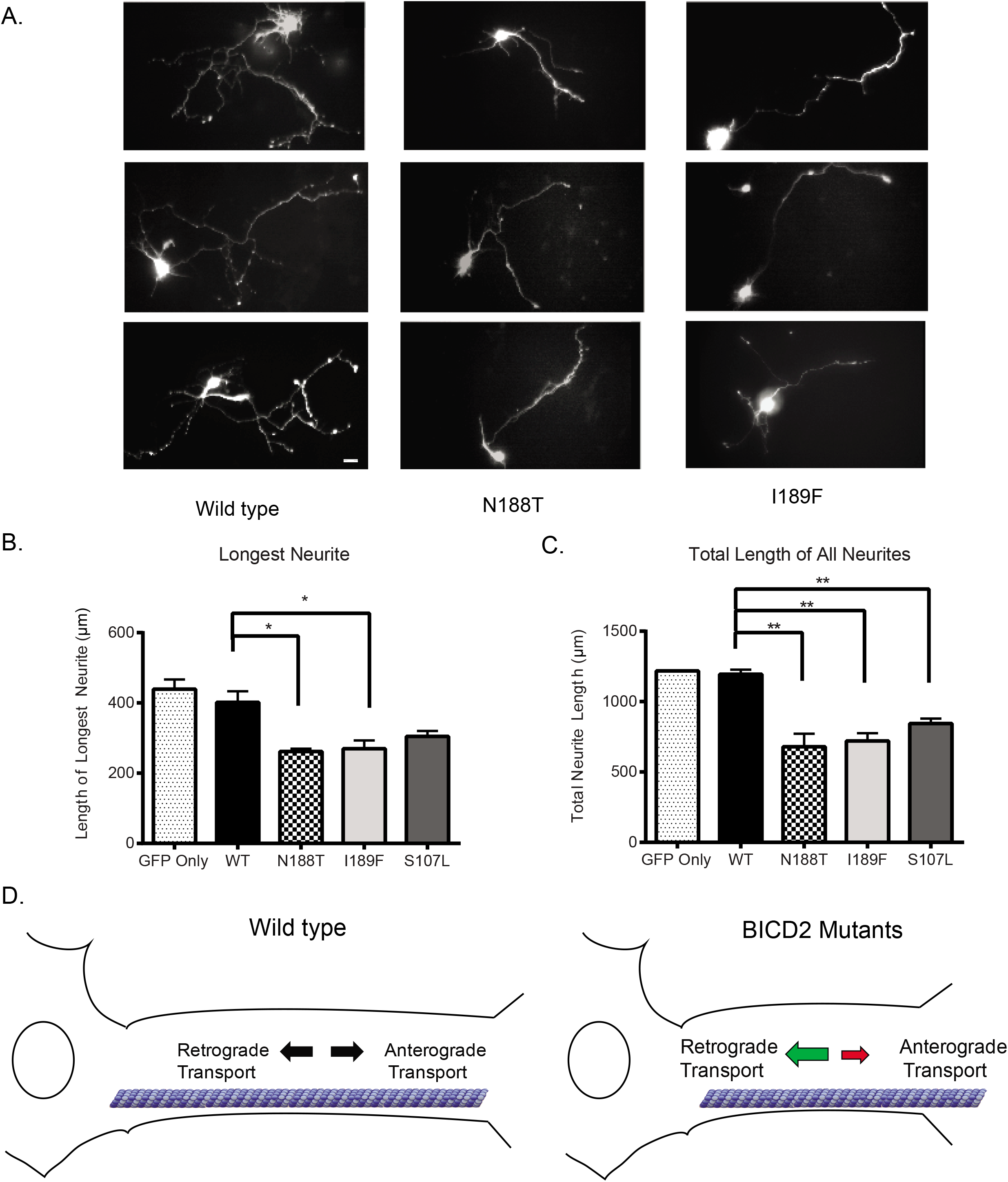
Over-expression of BICD2 mutants in rat hippocampal neurons results in a decrease in neurite length. **(A)** Representative images of hippocampal neurons over-expressing mCherry-BICD2_FL_ (wildtype and mutants) or an empty plasmid control; neurons were also co-transfected with soluble GFP in order to mark the neurites. Scale Bar: 10 μm. **(B)** Axon length after 3 days of overexpression of BICD2_FL_ constructs; mean and SD from n = 3 independent experiments (each experiment measuring 15-20 neurons for each construct) [*P ≤ 0.05]. **(C)** Total neurite length after 3 days of over-expression of BICD2_FL_ constructs; mean and SD from n = 3 independent experiment (each experiment measuring 15-20 neurons for each construct) [**P ≤ 0.01]. **(D)** Model of how a gain-of-function in dynein-based motility from BICD2 mutants might cause an imbalance of axonal transport and lead to SMA. See Discussion for more details.

In conclusion, our vitro and cell-based assays recapitulate Rab6a^GTP^-dependent activation of DDB motility and reveal that three BICD2 disease-associated mutations and one Drosophila mutant all display a similar phenotype of eliciting more dynein-dynactin motility. These effects were not evident with the Rab6a-independent, constitutively active BICD2_25-400_ construct. These results suggest a scenario in which the mutants lower the energy barrier for Rab6a^GTP^ to convert the autoinhibited BICD2_FL_ into an active conformation. Future work can test this possibility by using assays that probe the conformational states of BICD2_FL_.

Previous cell-based models have suggested that BICD2 disease mutants may have a dominant negative effect on dynein function resulting in disrupted Golgi morphology (Burkhardt et al., 1997; Harada et al., 1998; Neveling et al., 2013). While we cannot rule out dominant negative effects that cannot be measured with our assays, our data suggest that the mutants produce a dominant positive effect leading to enhanced dynein cargo transport. While subtle (∼25% increase in motile DDB complexes), such effects may perturb transport in an axon (which can be up to 1 meter long in a motor neuron) over time.

To explain the SMA-LED2 disease phenotype, we propose a model in which mutant BICD2 increases dynein activation and leads to an imbalance between anterograde and retrograde transport (Figure 5D). The shift towards dynein transport could produce a decrease in the delivery of cargo to the nerve terminal, which in turn could lead to a gradual decline in function of neurons with long axons. This model is congruous with experimental data from transgenic mouse models. BICD2 null mice possess a normal spinal cord and their motor neurons exhibit no evidence of abnormal retrograde transport (Jaarsma et al., 2014). These mice, which die at around p30, do not display a SMA phenotype, suggesting that the absence or inhibition of BICD2 activity cannot explain the disease etiology. In contrast, transgenic mice expressing the shorter, more active N-terminal construct of BICD2 were impaired in retrograde axonal transport and Golgi fragmentation, presumably through a dominant negative effect of activating dynein and dynactin without cargo and causing its accumulation in the cell body (Teuling et al., 2008). Curiously, however, these mice, which live up to two years, also did not develop any motor abnormalities. The phenotypic differences in mice expressing the N-terminal construct of BICD2 compared with haploid S107L, N188T, and I189F mutations suggests that the C-terminus of the BICD2 protein that interacts with cargo is necessary for contributing to the SMA disease phenotype.

Recently, Hoang et al. (2017) showed that disease-related mutants in the human dynein heavy chain, including three mutations associated with SMA, exhibited normal DDB complex formation and velocity, but displayed a ∼60-70% reduction in processivity. These results differ from those described for the SMA BICD2 mutants, which show no change in processivity and a gain-of-function increase in the amount of retrograde transport. Taken together, these studies on dynein and BICD2 mutations reveal multiple ways in which changes in motor activity can affect can affect the homeostasis and function of neurons in SMA.

## Acknowledgments

We thank N. Stuurman for help with microscopy, and R. McKenney for both technical help and discussions, and C. Schroeder, Y. Wang and S. Niekamp for discussions as well. This work was supported by a MIRA grant from NIGMS (NIH 1R35 GM118106).

### Author Contributions

W. Huynh performed the experiments. W. Huynh and R. Vale designed the experiments and wrote the manuscript.

## Materials and Methods

### DNA constructs

The cDNAs for mouse BICD2 (accession #AJ250106.1) and mouse Rab6a (accession #BC019118.1) were obtained from the Thermo Scientific MGC collection. BICD2 N-terminal constructs were cloned into a pet28a vector containing an N-terminal 6xHis-strepII-sfGFP tag. Full-length BICD2 WT and mutant constructs were cloned into a pFastbac vector containing either an N-terminal StrepII-sfGFP or StrepII-SNAPf tag followed by a C-terminal 6xHis tag for tandem purification. Rab6a was cloned into a pGEX vector containing an N-terminal GST tag followed by a PreScission protease cleavage site. The Rab6a constructs span the GTPase domain of Rab6a (a.a. 8-195), followed by a short linker and either a KCK motif for maleimide labeling purposes (liposome experiment), or a His10-tag. Point mutations for both the BICD2 constructs and the Rab6a GDP/GTP (Q72L and T27N, respectively) constructs were created by the protocol of Phusion, Inc.

For the cell-based peroxisome assay, the sequence spanning amino acids 1-42 of human PEX3 was synthesized via a gBlock (IDT), and was cloned into the N-terminal of a modified eGFP vector, followed by a C-terminal FRB sequence. BICD2 constructs for this assay were cloned into a pHR vector containing an N-terminal mCherry sequence and C-terminal FKBPf36v sequence. The Rab6a-PEX construct was constructed by fusing the PEX3 sequence to the C-terminus of Rab6a lacking the cysteines for lipidation (1-203aa). A mTagBFP2 was fused to the N-terminus of this construct as well. For neuron expression, BICD2 was cloned into an N-terminal pmCherry vector.

### Protein Purification

BICD2 N-terminal and Rab6a constructs were transformed into the *Escherichia coli* strain BL21 RIPL from Agilent. Cells were grown in LB at 37°C until growth reached ∼0.6 OD_280_. The temperature was then lowered to 18°C and cells were induced over-night with 0.5 mM IPTG. Bacterial pellets were resuspended in either strep lysis buffer (30 mM HEPES, pH 7.4, 300 mM NaCl, 2 mM MgCl_2_, 5% glycerol, 5 mM DTT, 1 mM PMSF) for BICD2 or PBS containing 5% glycerol, 5 mM DTT, and 1 mM PMSF for Rab6a. Cells were lysed using an Emulsiflex press (Avestin) and clarified at 40,000g for 60 min. Lysates were bound to either Strep-tactin Superflow Plus resin (QIAGEN) or GST Sepharose 4B (GE Lifesciences). Beads were then washed extensively and the bound protein was eluted with 3 mM desthiobiotin in lysis buffer (BICD2) or via overnight cleavage with PreScission (Rab6a).

BICD2 full-length constructs were purified from SF9 cells and purified using tandem purification. Bacmids isolated from DH10Bac cells were transfected into SF9 cells. P2 virus was used to infect SF9 cells grown in shaker flasks to a density of 2x10^6^ cells/ml. After ∼60 hr of infection, cells were harvested, pelleted, and resuspended in NiNTA lysis buffer (30 mM HEPES pH 7.4, 300 mM NaCl, 5% glycerol, 20 mM imidazole, 1 mM PMSF, 5 mM βME) and lysed using an Emulsiflex. The lysate was clarified at 40,000g for 1 hr and bound to NiNTA resin from Qiagen. After extensively washing with lysis buffer, cells were eluted with lysis buffer containing 500 mM imidazole. The eluted material was then diluted to lower the imidazole concentration to 100 mM, and bound to Streptactin beads as described above. After elution, proteins were concentrated and injected onto either a Superose 6 10/300GL column (BICD2 constructs) or a S200 10/300GL column (Rab6a) from GE healthcare. Gel filtration buffer was comprised of: 30mM HEPES pH 7.4, 150mM NaCl, 10% glycerol, 2mM TCEP. Peak fractions were pooled and concentrated and then flash frozen. For SNAP-fusion constructs, purified proteins were bound to Streptactin beads and incubated at a 2:1 ratio of SNAP-TMR Star:Protein on ice for 1 hour. Beads were briefly washed in the gel filtration buffer and the protein was eluted with 3mM of desthiobiotin in gel filtration buffer.

Dynein and dynactin were prepared from RPE-1 cells. Fifteen 15cm dishes of RPE-1 cells were grown to ∼80-900% confluency and then washed with PBS. The cells were then harvested using a cells scraper and pelleted at 500 x g for 5 minutes. They were then resuspended in 2.5mL of Buffer A (30 mM HEPES pH 7.4, 50 mM potassium acetate, 2 mM magnesium acetate, 1 mM EGTA, 10% glycerol) along with 5mM DTT, 1mM PMSF, and 1% Triton X-100 and incubated on ice for 10 minutes. The lysate was clarified at 266,000 x g and diluted to a volume of 25mL using Buffer A containing 1mM PMSF and 5mM DTT to reach a final Triton X-100 concentration of 0.1%. Streptactin sepharose (GE Healthcare) bound to purified Strepll-SNAP-FIP3 was added to the lysate and incubated for ∼1 hour. The beads were then washed 4x with Buffer A containing 5mM DTT and 0.1% NP-40. Buffer A containing an additional 300mM NaCl was then added to the beads to form a 50% slurry and incubated on ice for 10 minutes to elute the dynein and dynactin bound to FIP3. This slurry was then passed through a 0.22μm filter to remove the beads. An equal volume of 50% Streptactin sepharose was added to this elution and allowed to gently mix for 30 minutes at 4°C to remove any remaining Strepll-SNAP-FIP3. The slurry was then filtered again and then diluted with Buffer A to reach a final NaCl concentration of 200mM. Sucrose was added to a final concentration of 6% and the dynein/dynactin were then flash frozen as aliquots.

### Porcine Brain Pull-downs

Pull-downs were performed as previously described (McKenney et al., 2014). 500μl of clarified porcine brain lysate in Buffer A (30 mM HEPES pH 7.4, 50 mM potassium acetate, 2 mM magnesium acetate, 1 mM EGTA, 10% glycerol) was added to 80μl of a 50% suspension of Streptactin Sepharose beads (GE Healthcare) along with 100 nM of the BICD2 construct to be tested. To this, 0.1% of NP-40, 1 mM PMSF, and 5 mM DTT were added and the mixture was incubated at 4°C for 1 hr. The beads were then pelleted and washed 4 times with Buffer A containing 0.1% NP-40 and 5 mM DTT. The proteins were then eluted using SDS loading buffer and resolved on an acrylamide gel (Invitrogen). For experiments where Rab6a was supplemented, the appropriate amount of recombinant Rab6a protein was added prior to the 4°C incubation step and processed in the same fashion.

### Immunoblot Analysis

The SDS-PAGE gel was transferred onto a nitrocellulose membrane using the iBlot system from Invitrogen. Western blotting was performed using TBS buffer with 0.1% Tween-20 (TBS-T). Membranes were blot for 1 hr at room temperature with 5% milk in TBS-T, followed by a 1 hr primary antibody incubation. Antibodies used are as follows: mouse anti-p150 (clone 12, 1:250; BD), mouse anti-dynein intermediate chain (clone 74.1, 1:1000; EMD Millipore), mouse anti-6xHis (1:1000, Roche), mouse anti-Rab6a (ab55660, 1:500, Abcam). Membranes were washed 3x with TBS-T and then incubated with a secondary antibody for 1 hr at r.t. Secondary antibodies were: anti-mouse-800 or anti-rabbit-680 (1:10,000; Molecular Probes). Membranes were washed 3x with TBS-T and then imaged using an Odyssey Clx Infrared Imaging System (LI-COR Biosciences). Quantification of band intensity was performed using ImageJ’s (NIH) gel analysis feature. P values were calculated using a one sample *t* test with a hypothetical mean of 1.

### Liposome preparation and Rab6a labeling

Liposomes were prepared from a mixture of lipids dissolved in chloroform at (82.4% POPC, 15% POPS, 2.5% 18:1 PE MCC, 0.1% rhodamine PE (Avanti)). After mixing, they were dried under a constant stream of argon and then desiccated in vacuum overnight. The lipid film was then resuspended with Buffer A without glycerol and passed 25X through an extruder containing a 200 nm pore size polycarbonate filter. A 3:1 ratio of Rab6a to maleimide lipid was then mixed together and incubated overnight. The next day, DTT was added to a final concentration of 10 mM to quench the reaction, and the liposomes were pelleted at 160,000g for 20 minutes to remove unlabeled Rab6a. The pellet was then washed and resuspended with Buffer A.

### Single-molecule and liposome imaging

#### Preparation of microtubules

Microtubules were prepared as previously described (McKenney et al., 2014). Unlabeled tubulin was mixed with biotinylated tubulin and Alexa-640 labeled tubulin at a ratio of ∼10:2:2 in BRB80 (80 mM PIPES, pH 6.8, 1 mM EGTA, 1 mM MgCh). 5 mM GTP was added and the mixture was incubated in a 37°C water bath for 10 min before adding. 20 μM of Taxol and incubated for an additional 50 min at 37°C. Microtubules were spun over a 25% sucrose cushion at 160,000 g for 10 min in a tabletop centrifuge prior to use.

#### Preparation of DDB complexes and imaging

DDB complexes were formed in a 30 μL reaction volume containing ∼20 nM of recombinant BICD2, ∼0.15 mg/ml of dynein/dynactin, and Buffer A along with 0.1 mg/ml Biotin-BSA, 0.5% pluronic acid F-127, and 0.2 mg/ml κ-casein (Buffer B). The proteins were mixed and incubate on ice for 20 min prior to use. In experiments where Rab6a was also included, the respective construct was added to a final concentration of 1 μM. Flow chambers with attached microtubules were performed as described (Schroeder and Vale, 2016). A 1:4 or 1:5 dilution of the DDB complex in Buffer B was then added in presence of 1 mM ATP and the Trolox/PCA/PCD oxygen scavenging system (Aitken et al., 2008). For processivity experiments, a SNAP-BICD2 construct labeled with the TMR-Star fluorophore from NEB was used. Because the run-lengths using Buffer B tended to span the entire lengths of our microtubules, we added 15 mM of KCl to Buffer B (termed Buffer C) for measurement of processivity. The increased amount of salt in Buffer C reduced the run length for DDB (∼4 μm) so that it could be measured more accurately on microtubules (average length of ∼16 μm used in these measurements).

For liposome assays, the same components as in the DDB mixture above was mixed in a tube, along with either Rab6a^GTP^ or Rab6a^GDP^ liposomes. The liposomes were diluted to a final concentration of ∼2.5 μM total lipid. After incubation, the solution was flowed in undiluted into an imaging chamber after addition of 1 mM ATP and Trolox/PCA/PCD. The amount of BICD2-GFP fluorescence on the liposomes was used to determine the average number of molecules bound to each liposome by comparing it to that of single GFP molecules captured via biotin-anti-GFP stuck onto a coverslip coated with streptavidin. Fluorescent intensities were measured by integrating the average fluorescence intensity over 5 frames at 200ms exposure and quantified with the FiJI plugin Spot Intensity Analysis (http://imagej.net/Spot Intensity Analysis). Liposome GFP fluorescence (bound to glass) was normalized to single GFP fluorescent yielded an average of 6.01 BICD2 molecules per Rab6a^GTP^ liposomes (n = 6000 intensity measurements of single GFP and n = 728 intensity measurement of BICD2-GFP on liposomes).

Total internal reflection fluorescence images were acquired on a Nikon Eclipse TE200-E microscope equipped with an Andor iXon EM CCD camera, a 100x 1.49 NA objective, and MicroManager software (Edelstein et al., 2010). Exposures of 100 or 200 ms with a frame interval of 1 per sec for 300 sec was typically utilized for the acquisition. All assays were performed at room temperature.

### Peroxisome Assay in U2OS cells

U2OS cells from ATCC were cultured in DMEM medium containing 10% FCS and 1x penicillin/streptomycin/glutamine. Cells were seeded in glass-bottom 96 well plates and were at ∼40% confluence on the day of transfection. Cells were transfected with TransIT Lt1 as per the manufacturer’s protocol with a combination of the PEX-eGFP-FRB, mCherry-BICD2-FKBP, and PEX-Rab6a plasmids and incubated overnight. The next day, 1 μM of rapamycin or DMSO were added to cells and allowed to incubate at 37°C for 1 hr. 10 min before the end of this incubation, Cell Mask Far Red (Thermofisher) was added (0.5x of the manufacturer’s solution) to stain the plasma membrane. Cells were then washed with PBS and fixed with 4% paraformaldehyde (PFA) at room temperature followed by extensive washing with PBS. Cells were imaged at room temperature using an inverted Nikon Eclipse Ti microscope using a 0.75 NA 40x air objective, Andor Xyla camera, and Micro-Manager software (Edelstein et al., 2010).

### Primary hippocampal neuron cultures and transfection

Primary hippocampal neuron cultures were prepared from tissue from embryonic day 18 (E18) rat brains which were shipped over-night on ice from Brainbits LLC. Cells were prepared according to provider’s instructions. Briefly, the tissue was incubated with a 2 mg/ml solution of papain in Hibernate E without calcium (Braintbits) for 10 min at 30°C. The tissue was then triturated using a fire polished Pasteur pipette for 1 min and then large tissue pieces were allowed to settle for 1 min. The supernatant was collected and centrifuged at 200xg for 1 min to collect cells. After removal of most of the supernatant, the pellet was resuspended with 1 ml of NbActiv1 and the cells were counted after staining with Trypan Blue (1:5 dilution). ∼50,000 cells were seeded onto poly-d lysine/laminin coated round coverslips (Corning BioCoat) placed into wells of a 24-well plate.

One day after plating, neurons were transfected using Lipofectamine 2000 (Invitrogen). Each transfection reaction contained 1 μg of total plasmid DNA along with 4 μL of Lipofectamine reagent incubated together for 20 min at r.t. Media from the neurons was then removed and saved and fresh NbActiv was added to the wells along with the transfection mixture. After 2.5 hr at 37°C in 5% CO2, the neurons were washed with NbActiv and the original conditioned media was added back. Cells were fixed 3 days after transfection using 4% PFA for 20 min at room temperature. After washing, the coverslips were mounted onto slides using Vectashield (Vector Labs). Neurons were imaged for both soluble GFP and mCherry-BICD2 signal at room temperature using a Zeiss Axiovert 200M microscope with a 0.5 NA 20x air objective, Hamamatsu C4742-98 CCD camera, and Micro-manager software (Edelstein et al., 2010).

### Image Analysis and Quantification

#### Analysis of DDB and liposome motility and run length

Kymographs were created from the single molecule or liposome movies using ImageJ. The number of runs per micron of MT per unit of time was obtained from these kymographs, with each run being scored if it was >1 μm in length. P-values were calculated using an unpaired *t* test. The cumulative frequency was used for analysis of run lengths, as previously described (McKenney et al., 2014). The 1-cumulative frequency distribution was fitted with a one-phase exponential decay for run lengths of greater than 1uM, as runs shorter than this were undersampled or difficult to measure given our imaging conditions (Thorn et al., 2000).

#### Analysis of peroxisome clustering

The images acquired from fixed U2OS cells were processed and analyzed using the Compaction Profiler plugin previously developed for the ICY software (Mounier et al., 2012). Briefly, the fluorescent signal from the GFP channel corresponding to peroxisomes were detected after masking the shape of cell via the CellMask channel and quantified using the Spot Detector plugin in ICY. The detected fluorescence data was then passed along to the Compaction Profiler plugin, which then calculated the area of an ROI inside the cell that represents 90% of the total fluorescence signal. A higher degree of clustering results in a smaller calculated area, whereas a more diffusive signal yields a larger area. P-values were calculated using an unpaired *t* test.

#### Analysis of primary hippocampal neurons

Neurite lengths were measured using the soluble GFP signal from the transfected plasmid. Acquired images were processed via ImageJ. The background was subtracted using the default setting, followed by a median filter with a value of 2.0. The images were adjusted for brightness and contrast and the ImageJ plugin NeuronJ was then utilized to trace all neurites including branches from individual neurite. The longest neurite from a cell was also identified and measured. P-values were calculated using an unpaired *t* test.

## Supplemental Materials Summary

Fig. S1 shows a comparison of a brain lysate pull-down between BICD2_25-400_ and BICD2_FL_, a Rab6a blot for the titration experiment in Fig. 1 along with the quantification of DIC and p150 intensity for Rab6a^GTP^, single molecule quantification of BICD2_25-400_ with or without Rab6a, and a GST pull-down showing Rab6a binding to BICD2_FL_ but not the truncated BICD2. Fig. S2 shows quantification of the pull-down experiment in Fig. 2, processivity and motility with Rab6a^GDP^ between WT and mutants, a pull-down using the N-terminal construct of either WT and mutant, and velocity of liposomes from Fig. 3. Fig. S3 displays peroxisome experiments in which Rab6a-PEX is co-transfected as well as neuron data for the fly mutant and an analysis of the number of processes emanating from the neurons for both WT and mutants.

**Fig. S1.**
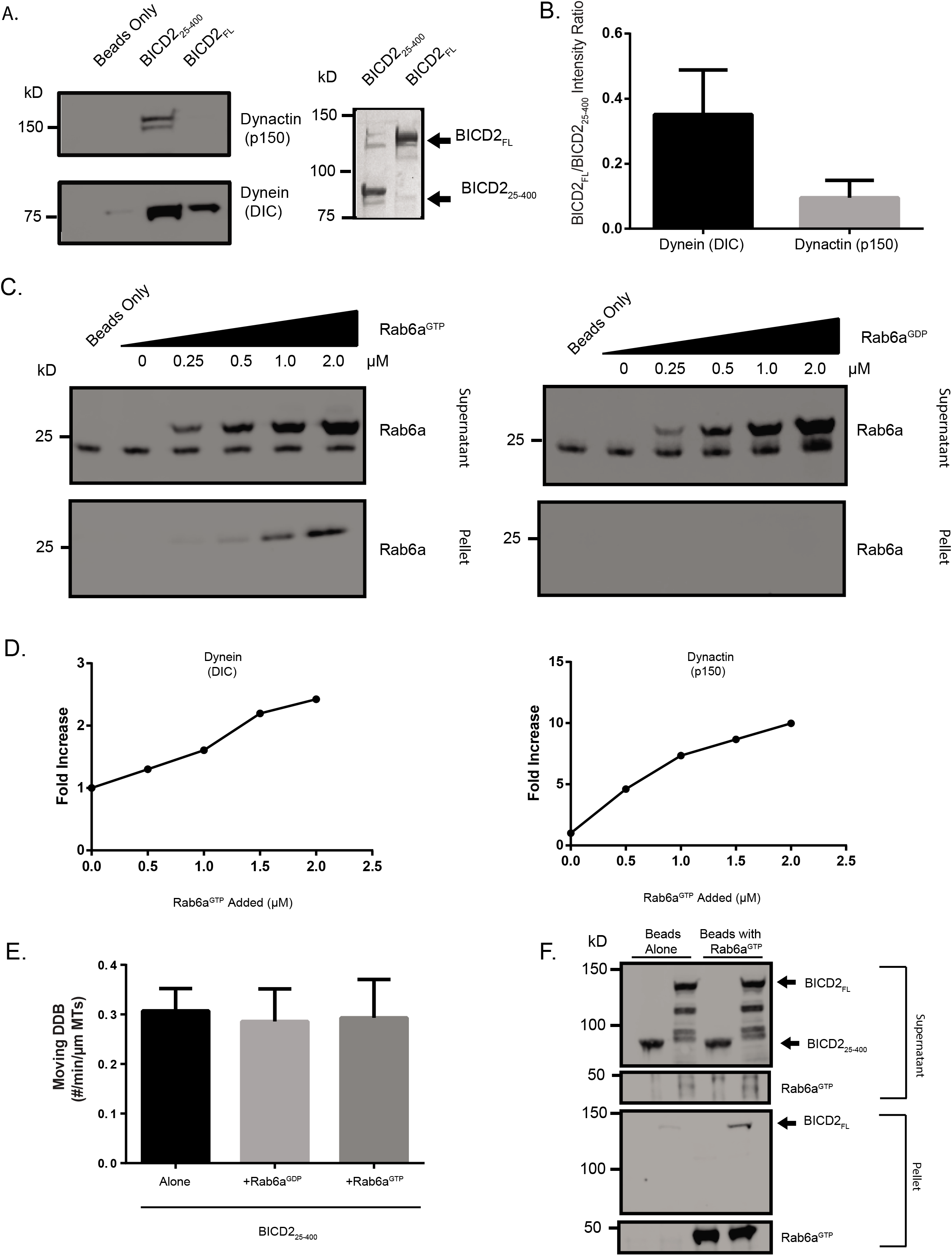
BICD2_25-400_ recruits more dynein as compared to full-length. **(A)** A comparison of the amounts of dynein and dynactin pulled down from porcine brain lysate between BICD2_25-400_ and BICD2_FL_ **(B)** Quantification of the pull-down; mean and SD from n = 3 independent experiments. **(C)** The supernatant and pellet fraction of the pull-downs from Figure 1B showing that Rab6a^GTP^ but not Rab6a^GDP^ binds to the BICD2 on beads. **(D)** Quantification of the western blot shown in Figure 1B for the case of Rab6a^GTP^ addition. Values are normalized to the lane in which no Rab6a is added. **(E)** Quantification of number of moving motors per micron of microtubule of BICD2_25-400_ with either no Rab6a added, Rab6a^GDP^ added, or Rab6a^GTP^ added; mean and SEM from n = 3 independent experiments (each experiment measuring a minimum of 30 microtubules). **(F)** GST-Rab6aGTP pull-down of either BICD2_25-400_ or BICD2_FL_.

**Fig. S2.**
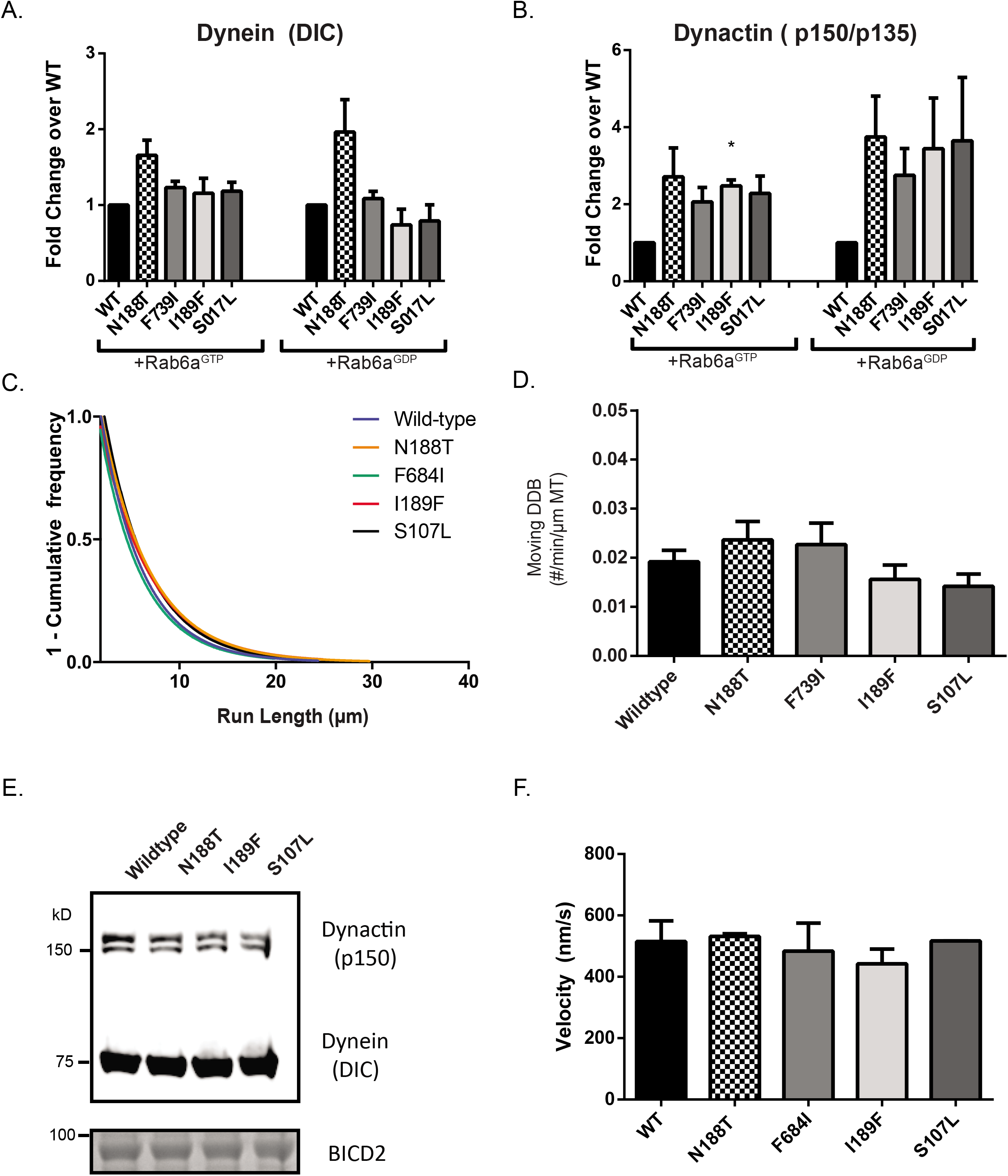
N-terminal mutant constructs of BICD2 are comparable to WT. **(A)** and **(B)** show the quantification of the western blots for Rab6a^GTP^ and Rab6a^GDP^ pull-downs from Figure 2; mean and SEM from n = 3 independent experiments. For (A), GTP addition mean and SEM are: N188T: 1.66 ± 0.20, p=0.082; F739I: 1.23 ± 0.082, p=0.11; I189F: 1.16 ± 0.20, p=0.51; S107L: 1.18 ± 0.12, p=0.26. GDP: addition N188T: 1.96 ± 0.42, p=0.15; F739I: 1.08 ± 0.097, p=0.47; I189F: 0.738 ± 0.21, p=0.34; S107L: 0.792 ± 0.21, p=0.43. For (B), GTP addition mean and SEM are: N188T: 2.71 ± 0.75, p=0.15; F739I: 2.06 ± 0.375, p=0.11; I189F: 2.48 ± 0.15, p=0.011; S107L: 2.28 ± 0.45, p=0.10. GDP: addition N188T: 3.75 ± 1.06, p=0.152 F739I: 2.75 ± 0.69, p=0.13; I189F: 3.44 ± 1.3, p=0.20; S107L: 3.65 ± 1.6, p=0.25. (C) Processivity data in the form of a 1-cumulative frequency histogram for one replicate, comparing different full-length BICD2 constructs. (D) The motility assay for DDB + Rab6a^GDP^ is also shown; mean and SEM from n = 3 independent experiments. (E) A comparison of BICD2_25-400_ for WT and appropriate mutant constructs in the porcine brain lysate pull-down assay. Shown are the western blots of the pellet fractions for p150 and DIC and a Coomassie staining for BICD2. (F) Velocity measurements of liposomes shown in Figure 4D; mean and SEM from n = 3 independent experiments. Each experiment measured >100 liposomes. The error bar for S107L is too small to appear on the graph.

**Fig. S3.**
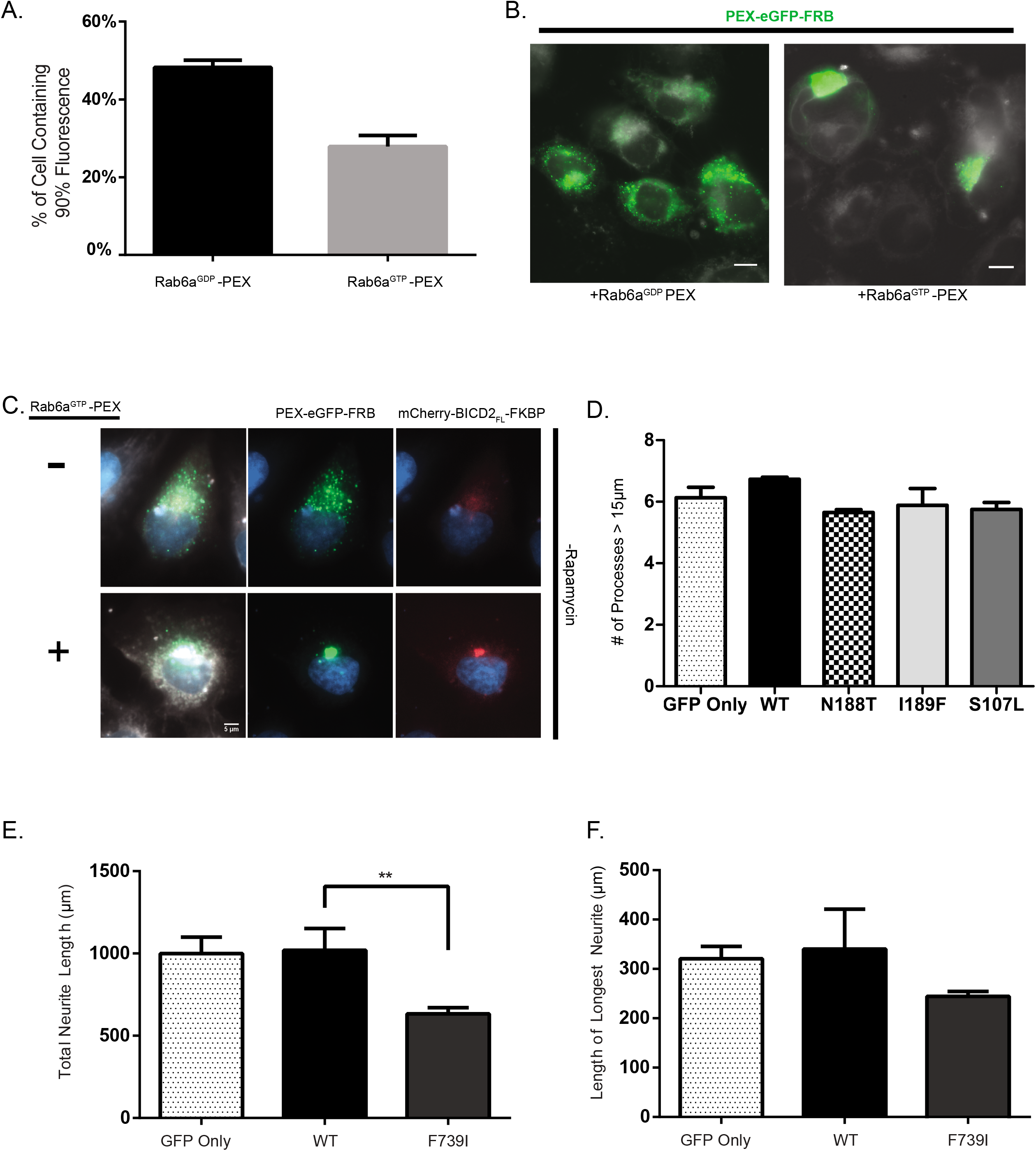
Peroxisome-localized Rab6a^GTP^ is sufficient for clustering in cells. **(A)** A comparison of peroxisome clustering when either the GDP or GTP of Rab6a is fused with the PEX sequence; mean and SEM from n = 2 independent experiments. Each experiment measured at least 30 cells. **(B)** Representative images of peroxisomes from (A). The green color reflects PEX-eGFP-FRB, while the white is from a membrane dye. Scale Bar: 10 μm **(C)** Example of BICD2_FL_ localizing to the peroxisomes even in the absence of rapamycin. In the top row, cells were transfected with PEX-eGFP-FRB and mCherry-BICD2_FL_. In the bottom row, the cells were additionally transfected with Rab6a^GTP^-PEX as well. **(D)** Quantification of total number of processes (> 15 μm long) emanating from the cell body of the neurons from Figure 5a; mean and SD from n = 3 independent experiments. **(E)** Total neurite length after 3 days of overexpression of BICD2_FL_ constructs; mean and SD from n = 3 independent experiments (each experiment measuring 10-15 neurons for each construct) [**P ≤ 0.01]. **(F)** Axon length after 3 days of over-expression of BICD2_FL_ constructs; mean and SD from n = 3 independent experiments (each experiment measuring 10-15 neurons for each construct).

## Supplemental Movie Legends

**Video S1. Comparison of BICD2_25-400_ vs BICD2_FL_ DDB.** Movie shows motility of either BICD2_25-400_ (left) or BICD2_FL_ (right) DDB complexes (green) on microtubules (blue). Scale bar is 2 μm. See Figure 1C for details.

**Video S2. Comparison of Rab6a^GDP^ and Rab6a^GTP^ liposome motility.** Movie shows a comparison of Rab6aGDP (left) vs Rab6aGTP (right) liposomes (red) when incubated together with DDB_FL_. Microtubules are shown in blue. Scale bar is 2 μm.

